# A Meta-Analysis of 515 Human Proteomes Defines the Core Composition and Contaminants of Plasma Extracellular Vesicles isolates

**DOI:** 10.64898/2026.01.28.702342

**Authors:** Ana Sofia Carvalho, Juan M. Falcón-Pérez’, Hans C Beck, Rune Matthiesen

## Abstract

Plasma extracellular vesicles (EVs) are promising liquid biopsy biomarkers, yet their proteomic profiling remains confounded by co-isolated lipoproteins and soluble complexes. Here, we resolve this heterogeneity through a meta-analysis of 515 human plasma EV proteomes. We demonstrate that the reproducible core proteome is dominated by soluble host-response modules rather than unique vesicular markers. Deconvolution analysis resolved the isolate into distinct biological sources, distinguishing the hepatic secretome and immune complexes from vesicular signals. Crucially, canonical tetraspanins (CD9, ADAM10) clustered with platelet and erythrocyte proteins, indicating that the bulk tetraspanin signal derives from hematopoietic ectosome shedding. In contrast, ESCRT machinery (TSG101, ALIX) formed a distinct, lower-abundance cluster, physically separating endosomal biogenesis from membrane shedding. We provide a modular atlas that mathematically distinguishes the functional matrix of coisolates from vesicular cargo, establishing a new framework for interpreting biological origin in plasma EV proteomics.

## Introduction

Extracellular vesicles (EVs) have emerged as pivotal mediators of intercellular communication and promising sources of liquid biopsy biomarkers. While broadly defined by the International Society for Extracellular Vesicles (ISEV) as non-replicating, lipid-bilayer-delimited particles released by cells^1^, the practical isolation of EVs from complex biofluids yields a heterogeneous mixture of vesicles and coisolated non-vesicular (NV) contaminants, such as lipoproteins, protein aggregates, and cellular debris^2^. Consequently, ascribing specific functions or biomarkers to EVs requires a rigorous distinction between vesicular cargo and soluble impurities.

The composition of the core EV proteome remains a subject of intense debate, heavily influenced by the biological source and isolation methodology. In controlled cell culture environments, high-resolution studies have challenged historical assumptions. For instance, Jeppesen *et al*.^*3*^ utilized high-resolution density gradients to demonstrate that widely reported EV cargoes, such as DNA, histones, and Argonaute proteins, are largely associated with non-vesicular extracellular matter or amphisomes rather than classical exosomes. Similarly, Kugeratski *et al*.^*4*^ showed that the abundance of classical tetraspanins (CD9, CD63, CD81) varies significantly across cell lines, mirroring parental expression rather than a universal vesicle signature, and proposed Syntenin-1 as a more robust ubiquitous marker.

Translating these findings to human plasma, however, presents unique challenges due to the vast dynamic range and the presence of blood-specific contaminants. Recent large-scale proteomics efforts have attempted to define plasma-specific signatures. Hoshino *et al*.^*5*^ analyzed over 400 human samples to define extracellular vesicle and particle (EVP) proteomes, identifying tumor-derived signatures. Addressing the dynamic range, Wu *et al*^*6*^ employed magnetic bead capture (Mag-Net) to uncover signatures distinguishing neurodegenerative conditions. Most recently, Rai *et al*.^7^utilized density gradient separation to propose a hallmark panel of 182 proteins capable of distinguishing small EVs from non-EV particles.

Despite these advances, a consensus on the plasma EV proteome remains elusive. Studies comparing isolation methods, such as Takov *et al*.^*8*^, warn that methods yielding higher particle counts (e.g., precipitation or size-exclusion chromatography) often retain significant lipoprotein contamination (e.g., APOB) which can confound functional assays. Critically, it remains unclear to what extent proposed universal plasma EV markers are confounded by the co-isolation of highly abundant blood components, specifically those derived from platelet activation and red blood cell hemolysis.

Collectively, these studies underscore a critical unresolved question distinct from cell culture supernatants, what constitutes the vesicular proteome core of human plasma, and which reported markers are merely passengers of co-isolation? Whether defining a core proteome or establishing disease signatures, the assumed EV signature is inextricably linked to the rigor of separation and the presence of dominant contaminants like platelets and erythrocytes.

Therefore, in this manuscript, we aim to resolve this ambiguity through a large-scale meta-analysis of 515 human plasma EV proteomes. By integrating raw mass spectrometry data from diverse isolation methodologies with high-depth reference proteomes of purified platelets and erythrocytes, we deconvolve the plasma EV landscape into distinct variance components. We demonstrate that widely cited EV markers frequently segregate into clusters defined by hemolysis and platelet debris rather than vesicle biogenesis. Based on these findings, we establish robust core EV markers and source-specific contaminants, providing a standardized resource to improve the reproducibility of liquid biopsy studies.

## Results

Previous efforts on defining plasma EV signatures tend to start from a very positive assumption that the preparation is rich in EVs and consequently abundantly identified proteins by LC-MS would serve as a signature for EVs or at least be defined as EV associated proteins. The premise in the current study is that at least for plasma EV preparation there will be multiple components contributing to the final EV sample. This premise is supported by two observations: 1) plasma EV preparation is fundamentally different in overall protein composition compared to EVs from cell culture or other biofluids such as urine, pleural effusion or bronchoalveolar lavage^9^. 2) It is well accepted that MS-based proteomics of plasma samples identifies varying degrees of erythrocytes, platelets and blood coagulation proteins^10^. It is reasonable to assume that the same protein groups that affect plasma preparations will also plaque EV preparations. Additionally, for EVs a non-EV signature for lipoproteins^11^, coagulation^12^ and immune complexes^13^ are expected (Table 1 and Supplementary Data 1).

**Table 1.**
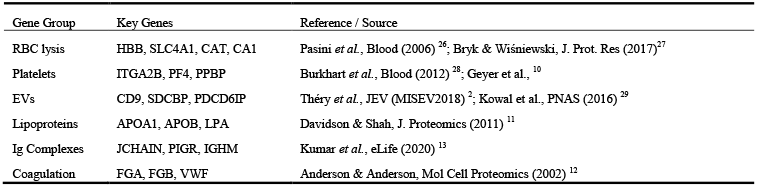
Summary of plasma EV anchor proteins.

### A Unified Proteomics Atlas of 515 Human Plasma EV Preparations

Data was restricted to label free quantification by DDA deposited in ProteomeExchange. Plasma EV data from ProteomeExchange were extracted from a total of 515 LC-MS files spanning across 20 publications (Fig. 1). In total 10 different EV isolation methods were applied across studies (Fig. 1a). DUC was the most frequent applied method across the studies and across LC-MS runs (Fig. 1a). The number of LC-MS runs per study ranged from 3 to 94 (Fig. 1b). The number of proteins identified per LC-MS run were for most studies stable with a few displaying variance across samples (Fig. 1c).

**Fig. 1.**
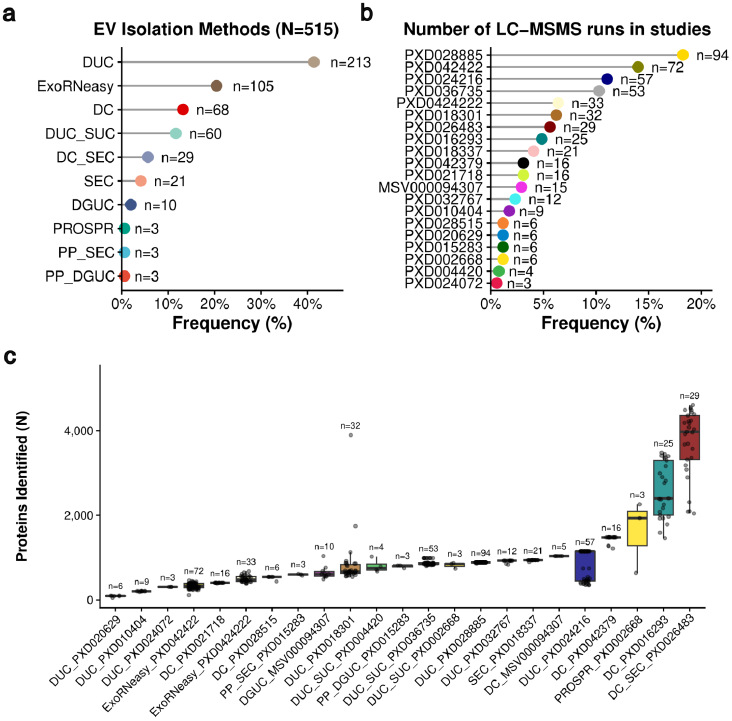
Overview of the 20 plasma EV proteomics studies. **a** Frequency of plasma EV isolation methods applied across 515 LC-MS runs. **b** Number of LC-MS runs per study. **c** Number of proteins identified for each EV isolation method and study. Note few studies used multiple isolation methods.

The range of proteins identified per study ranged from 142 to 5723 with a mean of 1552 and median of 1002. The majority of the identified proteins display a stochastic identification frequency. Meaning that these proteins were only identified in a few LC-MS runs (Fig. S1a and b). Fortunately there is a fairly large subset of 344 proteins that are consistently identified in more than 50% of all LC-MS runs (Fig. S1a and b insert). We evaluated the consistency of protein identification across independent LC-MS runs and diverse clinical cohorts to define the stability of the circulating EV proteome (Fig. S1). The data reveal a proteomics landscape characterized by a highly conserved core proteome amidst a backdrop of extensive biological variation. As expected for plasma, a biofluid that integrates vesicular signals from diverse tissues and pathological states, the majority of the proteome was detected stochastically, resulting in a right-skewed distribution of identification frequencies (Fig. S1a, c). This sparsity reflects the high dynamic range of the plasma proteome and the distinct biological signatures associated with the different pathologies included in this meta-analysis. Far from being technical noise, this variable proteome likely harbors the disease-specific biomarkers unique to each clinical condition. Remarkably, despite this systemic heterogeneity, we identified a robust core proteome present in the vast majority of samples (Fig. S1b). This ubiquitous subset likely represents the fundamental EV structural machinery alongside common co-isolated circulatory cargo, establishing a stable proteomics baseline independent of disease etiology. Notably, 25 of these core proteins are categorized as either EV components or common non-EV co-isolates by the MISEV2018 guidelines^2^. This includes six canonical EV markers: *HSPA8* (Hsc70), *SDCBP* (Syntenin-1), *ADAM10, FLOT1* (Flotillin-1), and the tetraspanins *CD9* and *CD81*^14^(Fig. 2a).

**Fig. 2.**
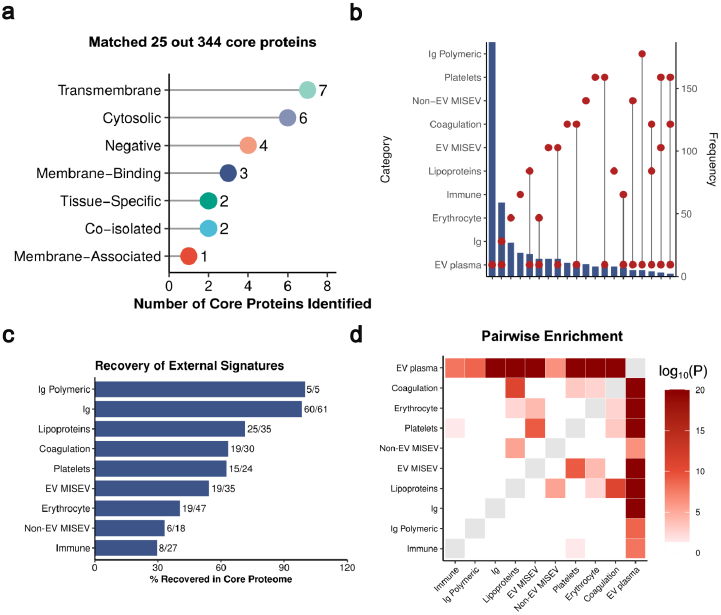
Overlap of Core Plasma EV Proteins with MISEV2018 Marker Categories and Contamination Signatures. **a** Number of core plasma EVs matching MISEV2018 marker categories. **b** Advanced Upset plot visualizing the overlap between our defined contamination signatures and core proteins. **c** Recovery plot demonstrating the overlap between our defined contamination signatures and core proteins. **d** Enrichment heatmap illustrating the degree of significant enrichment, with higher intensity represented by red. Self-comparisons (diagonal) are masked in gray to enhance contrast of the off-diagonal enrichment scores. Ig: soluble immunoglobulin.

A global heatmap of rank normalized iBAQ values for the core plasma EV proteome displayed strong clustering of individual LC-MS runs according to their study of origin. This indicates that technical variables, particularly the EV isolation strategy, significantly influence the quantitative abundance profiles of the core proteome, even among ubiquitously detected proteins.

In summary, a key finding is the consistent detection of 344 proteins across minimum 50% of all LC-MS runs, many of which exhibit similar high protein abundance (Fig. S2). This is encouraging, suggesting a degree of consistency across studies despite variations in pre-analytics, LC-MS sample preparation methods, and label-free LC-MS acquisition, as well as differences in the pathological state of the study subjects.

### Literature Review of Suspected EV and Non-EV Markers

Our analysis revealed a significant overlap between the core EV protein panel and established MISEV2018 marker categories. Specifically, 11 core proteins matched MISEV2018 EV membrane proteins and six cytosolic EV membrane proteins. This suggests a degree of alignment with established MISEV2018 EV markers and plasma EV core proteome (Fig. 2a). Despite this encouraging consistency, several non-EV markers were identified within the core proteome. Additionally, as mentioned above the premise of this study is that proteins identified in the core proteome of the plasma isolated EVs are from multiple biological sources. Therefore, literature mining for signatures from different biological sources was performed (Table 1 and Supplementary Data 1). To assess the potential contaminants within the core plasma EV proteomics data, we analyzed overlaps between our identified core plasma EV proteins and defined contamination signatures obtained by literature mining (Fig. 2). The advanced Upset plot (Fig. 2b) revealed multiple overlaps between potential contamination markers and the core plasma EV proteome. Suggesting that a substantial component of non-EV particles were isolated across studies. The recovery plot (Fig. 2c) further supported the presence of contamination signatures within the core plasma EV protein panel, demonstrating large overlap with for example immunoglobulins, lipoproteins, coagulation, platelets and erythrocytes. The enrichment heatmap (Fig. 2d) visually confirmed significant enrichment of our defined contamination signatures within the core plasma EV protein panel, with the highest levels of enrichment observed for MISEV2018 EV markers, soluble immunoglobulins, lipoproteins, coagulation, platelets and erythrocytes. In summary, these findings indicate that non-EV components contribute significantly to the core plasma EV proteome.

### Expression Level of Suspected EV and Non-EV Proteins

The previous section established that plasma EV proteomes contained proteins with markers originating for suspected EV contaminants and co-isolates. To further explore the severity of this contamination rank normalized iBAQ values were compared across contaminant markers and EV markers (Fig. 3a). It is clear that markers from possible EV contaminants are highly abundant in comparison to MISEV2018 EV markers. Also the contamination makers extracted from the literature in this study also display a higher abundance than the MISEV2018 non-EV markers. As expected MISEV2018 EV markers displayed a higher abundance than MISEV2018 non-EV markers (Fig. 3a). Overall coagulation, platelets, lipoprotein, soluble immunoglobulin and polymeric immunoglobulin markers display higher average abundance values than MISEV2018 EV markers (Fig. 3a). Note that the protein categories are defined as the core proteins together with proteins that are strongly associated with the category and these assignments are based on abundance values obtained after isolation of each of components corresponding to the category. For example, although abundant in platelets, FGB is synthesized by the liver and endocytosed by megakaryocytes (origin of platelets) from plasma^15^, justifying its classification as a plasma-associated rather than platelet-synthesized protein. Therefore, although FGB is a coagulation factor, its sequestration in alpha-granules means it is frequently associated with platelet contamination in EV preparations. Many of the core and strongly associated markers are shared among multiple signature categories, which complicate marker definition. Fig. 3a was created from a strict list of consensus markers. However, similar results of abundant non-EV protein signatures were obtained for a less strict definition of signatures (Fig. S3a). High average expression of non-EV proteins might be caused by a few studies displaying abnormally high contamination. To exclude this possibility the detectability across studies (Fig. 3b) and LC-MSMS runs (Fig. S3b) were calculated. Around half of the proteins in the signatures were identified in more than 75% of all studies and almost all in more than 50% of the studies (Fig. 3b).

**Fig. 3.**
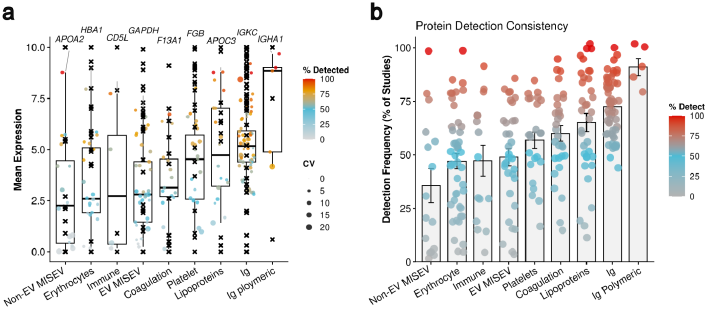
Quantitative abundance and detection consistency of core EV markers in comparison with non-EV markers. **a** Boxplots displaying the mean rank normalized iBAQ abundance of established EV markers (e.g., CD9, CD81) compared to common non-EV co-isolates (Lipoproteins, Erythrocyte proteins) across the entire dataset. The gene labels above each category indicate the most abundant protein in the core EV plasma proteome for the respective category. The central line represents the median, box limits indicate the 25th and 75th percentiles, and whiskers extend to 1.5* IQR. The crosses indicate abundance values from density gradient isolation of plasma EVs. **b** Detection frequency of each protein category, calculated as the percentage of the total studies in which the marker was identified. Note that non-EV proteins are both abundant (**a)** and exhibit high detection consistency compared to core EV markers (**b**).

Overall, non-EV proteins are abundant in plasma EV preparation and exhibit a high detection frequency across LC-MS runs and studies. Co-isolation of immunoglobulins and lipoproteins displayed in the highest abundance in the plasma EV preparations.

### The Reproducible Plasma EV Isolate: Co-Isolation of Functional Modules

To deconvolute the biological architecture of the plasma EV isolate, we restricted our analysis to the core proteome, defined as the 344 proteins consistently quantified in 50% of the 515 samples. This high-frequency dataset represents the stable, reproducible proteome background of clinical plasma EV preparations. We applied unsupervised hierarchical clustering to the rank-normalized abundances of these proteins to identify co-varying functional units (Fig. 4) and subsequently computationally validated these findings using a targeted candidate marker panel (Fig. 5).

**Fig. 4.**
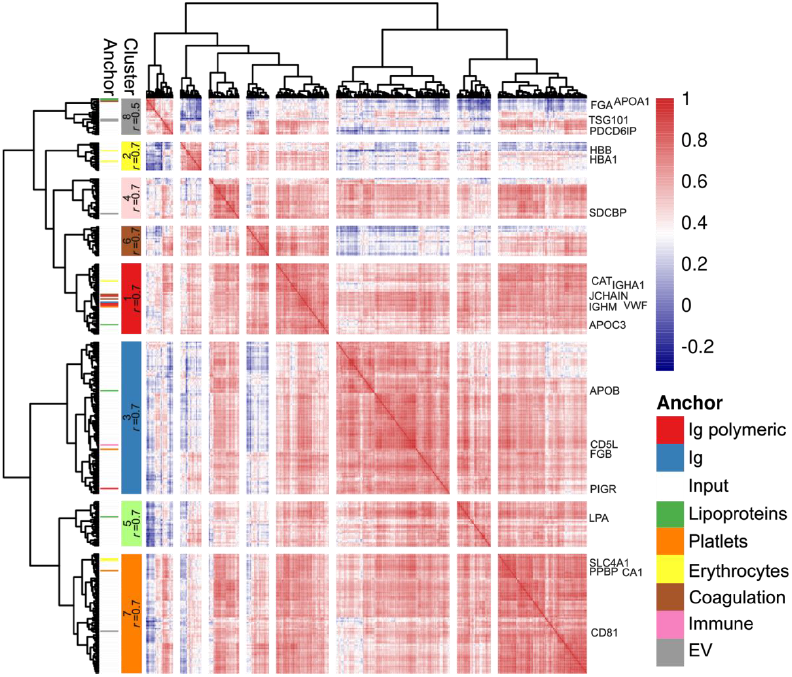
Resolution of the plasma EV core proteome into eight distinct biological modules. Pairwise correlation analysis of the core proteome reveals eight co-isolating protein clusters. Biological annotation was performed using the anchor proteins defined in Table 1. The side annotation bar indicates the specific biological category of individual proteins. Cluster-level color coding represents the predominant biological category identified within that module (determined by the most frequent anchor protein assignment); clusters lacking a dominant category are assigned a distinct color. Cluster labels indicate the Cluster ID and the mean correlation coefficient (r) of the module. Heatmap intensity represents the pairwise Pearson correlation between proteins.

**Fig. 5.**
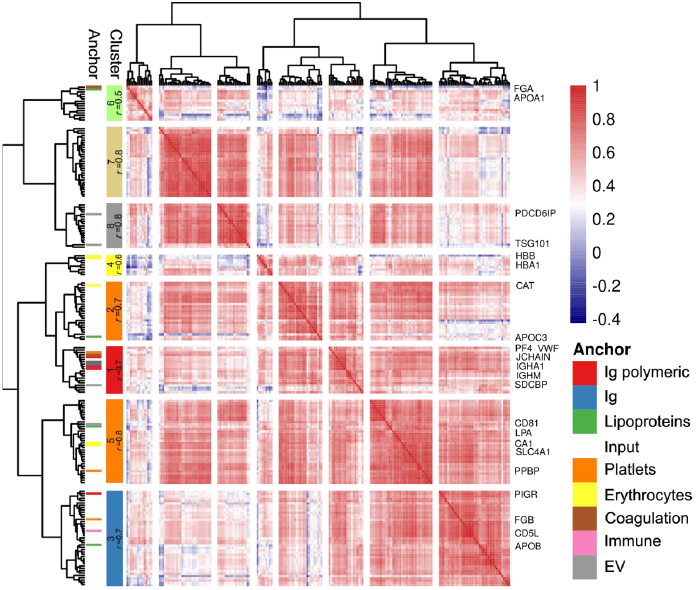
Application of the co-abundance framework to the Rai et al. candidate marker list. Unsupervised clustering of the candidate EV marker panel reported by Rai et al., performed using the same parameters as the core proteome analysis. Biological annotation and cluster color coding follow the exact scheme described in Fig. 4 and Table 1. The analysis resolves the consensus list into 8 distinct variance components, distinguishing vesicular markers from co-isolated plasma modules.

**Fig. 6.**
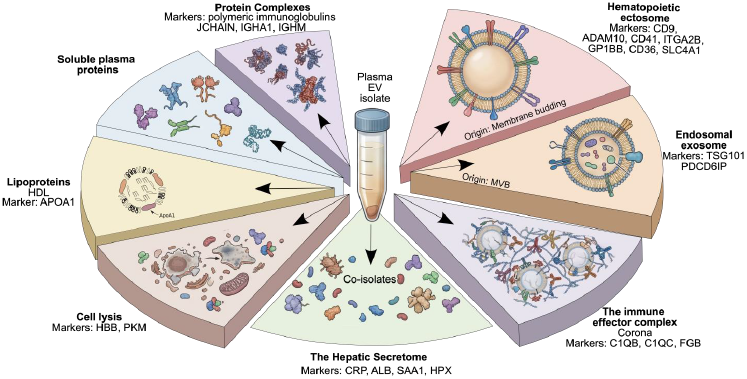
The modular architecture of the plasma EV isolate. The central schematic represents the bulk plasma EV isolate, often perceived as a homogeneous sample. The exploded sections illustrate the biological deconvolution of this isolate into distinct variance components, as resolved by our coabundance analysis. The Hematopoietic Ectosome module (Major Fraction) captures the bulk of the tetraspanin signal (CD9, ADAM10) and is derived from platelet/RBC membrane shedding. This is structurally distinct from the Endosomal Exosome module (Minor Fraction), which contains the ESCRT machinery (TSG101, ALIX). Additional slices represent the Functional Matrix of co-isolated contaminants, including the Immune Effector Complex (C1q, IgG), the Hepatic Secretome (CRP, Albumin), and Lipoproteins (ApoA1). Biomarker specificity requires resolving these independent biological sources rather than treating the isolate as a singular entity.

### The Core Proteome: Dominance of Soluble and Hematopoietic Backgrounds

Clustering analysis of the unbiased core proteome (Fig. 4 and Supplementary Data 3) revealed that the reproducible EV isolate is structurally dominated by non-vesicular plasma components and blood cell membrane fragments rather than generic tissue-derived vesicles.

The majority of the proteome segregated into high-abundance soluble modules driven by macromolecular size or surface adsorption. Cluster 1, Supplementary Data 3 *(r=0*.*72)* was enriched in polymeric immunoglobulins *(JCHAIN, IGHA1, IGHM)* and large glycoproteins *(VWF)*, representing massive molecular complexes that likely co-pellet due to their hydrodynamic radius. Similarly, Cluster 3, Supplementary Data 3 (*r=0*.*70*) captured the Immune Effector Complex, defined by the tight co-regulation of the C1 complement complex *(C1QB, C1QC)* and fibrinogen *(FGB)*. This cluster likely reflects the protein corona or immune complexes adsorbed to the vesicle surface.

Crucially, the primary cell-derived signature in the core proteome (Cluster 7, Supplementary Data 3, *r=0*.*74)* was a chimeric pan-hematopoietic ectosome module, driven by the co-isolation of platelet *(ITGA2B, GP1BB, CD36)* and erythrocyte *(SLC4A1)* membrane proteins. Definitive EV markers often cited in the literature, including CD9, CD81, and ADAM10, clustered inextricably within this hematopoietic module. This strongly suggests that the reproducible tetraspanin signal in plasma is driven by the bulk recovery of blood cell microvesicles rather than a generic exosome population.

Finally, the core analysis identified a specific endosomal/ESCRT signature in Cluster 8, Supplementary Data 3 *(r=0*.*54)*, anchored by the canonical markers TSG101 and ALIX (*PDCD6IP*). Notably, in this unbiased analysis, Apolipoprotein A-I (APOA1), the primary scaffold of HDL, also co-clustered with these markers. This indicates that while the ESCRT machinery forms a detectable variance component in the core proteome, its elution profile partially overlaps with High-Density Lipoproteins.

### Structural Resolution of Consensus EV Signatures

To benchmark our findings against the current consensus in the field, we applied our clustering framework to the extensive panel of EV features reported by *Rai et al?*. This dataset serves as a representative inventory of proteins frequently cited across the literature as plasma EV biomarkers. Our objective was to determine whether these widely reported features constitute a homogeneous vesicular signature or a composite of the broader plasma milieu. Unsupervised clustering of this representative feature set revealed that the canonical plasma EV signature is not a singular biological entity, but rather a composite of at least eight statistically distinct variance components (Fig. 5 and Supplementary Data 4). This analysis effectively disentangles the vesicular signals from the reproducible co-isolates that frequently confound plasma proteomics: 1) The hematopoietic ectosome core (Cluster 5 & 7). Consistent with our Core analysis, the strongest variance component in this consensus dataset (Cluster 5, *r=0*.*77)* defined a blood cell membrane signature. This module confirmed that canonical tetraspanins (CD9, CD81) and ADAM10, central pillars of EV definitions in the field, cluster inextricably with definitive platelet (ITGA2B) and RBC (SLC4A1) surface markers. This stratification clarifies a pervasive ambiguity in the field: the EV proteome signal in bulk plasma is primarily driven by the burden of hematopoietic ectosomes rather than generic tissue-derived exosome secretion. 2) Resolution of the endosomal vs. lipoprotein signal. A major strength of this structural analysis was the resolution of the ESCRT machinery from lipoprotein signals. While these two classes are often conflated in literature-derived lists, our clustering separated the endosomal markers TSG101 and ALIX (Cluster 8, *r=0*.*77)* from the soluble/micellar module containing Apolipoprotein A-I (Cluster 6). This demonstrates that while both vesicle types and HDL are present in consensus feature sets, they behave as independent biological variables that must be interpreted separately. 3) The misassignment of the soluble host response. Crucially, the clustering analysis revealed that a significant proportion of features broadly accepted as EV markers segregate into soluble plasma modules. Cluster 3 identified the Immune Effector Complex (C1q, Fibrinogen), while Cluster 1 captured soluble immunoglobulin complexes. The distinct segregation of these clusters from the membrane modules (Clusters 5 & 8) suggests that their frequent reporting in EV studies reflects their reproducible presence as part of the protein corona or soluble background, rather than vesicular encapsulation. In summary, by stratifying this representative dataset, we demonstrate that many proteins currently ascribed to the plasma EV proteome are actually markers of the soluble host response (immune complexes, coagulation) and hematopoietic turnover. While these non-vesicular coisolates may possess diagnostic utility as surrogates for systemic disease, distinguishing them from vesicular cargo is essential for accurately defining biological origin.

## Discussion

### Beyond Purity: A Modular Anatomy of the Plasma EV Isolate

The pursuit of a pure plasma EV signature has long been hindered by the complexity of the blood proteome, where vesicular signals are dwarfed by soluble proteins and lipoprotein particles. Our large-scale deconvolution of 515 clinical proteomes challenges the prevailing binary view of EVs vs. Contaminants. Instead, we demonstrate that the reproducible plasma EV isolate is a composite of at least eight distinct biological modules, each with a unique cellular origin and physical state. By resolving this architecture, we shift the analytical paradigm from simple inclusion lists to structural stratification. We show that high-abundance proteins frequently cited as EV markers do not behave as a homogeneous group, rather, they segregate into mutually exclusive components, ranging from the hepatic secretome^16^ and immune effector complexes to discrete hematopoietic ectosome and endosomal populations. This modular atlas explains why reproducibility in plasma proteomics has remained elusive in that different studies have been measuring the sum of independently regulated biological variables, liver function, immune activation, and platelet shedding, under the single label of EVs.

### The Divergent Biogenesis of Canonical Markers: Shedding vs. Sorting

A critical finding of this study is the definitive schism between the two primary modes of vesicle biogenesis in plasma. Current MISEV guidelines often group tetraspanins (CD9/CD81) and ESCRT machinery (TSG101/ALIX) as interchangeable markers of EV presence. Our data fundamentally overturns this assumption for plasma-derived samples.

We observed an inextricable clustering of CD9, CD81, and ADAM10 with the Platelet *(ITGA2B, GP1BB)* and Erythrocyte *(SLC4A1)* surface proteomes (Cluster 5, Table S4 and Cluster 7, Supplementary Data 3). This strongly implies that the vast majority of tetraspanin signal in plasma is not derived from generic tissue exosomes, but from the bulk shedding of hematopoietic ectosomes (microvesicles). This aligns with recent flow cytometry evidence suggesting that platelet-derived EVs constitute the dominant vesicular fraction in blood^17,18^. Consequently, we caution that elevations in CD9 or ADAM10 in liquid biopsies should be primarily interpreted as indices of hematopoietic activation or turnover, rather than tumor burden.

In contrast, the canonical ESCRT machinery (TSG101, ALIX) formed a statistically distinct, lower-abundance module (Cluster 8, Supplementary Data 3 and 4). This separation suggests that endosomal exosomes represent a minor fraction of the total isolate, chemically and physically distinct from the dominant ectosome background. This mirrors findings from high-resolution iodixanol gradient studies^3^, which demonstrated the physical separation of tetraspanin-rich and ESCRT-rich vesicles. Our analysis confirms this dichotomy exists even in the complex milieu of clinical plasma.

### The Functional Matrix: Redefining Contamination as Functional Co-Isolation

Our analysis revealed that the core proteome is dominated by soluble modules that are chemically distinct from vesicles yet reproducibly co-isolated. Rather than dismissing these as random noise, we characterize them as the functional matrix of the plasma isolate.

Specifically, Cluster 3, Supplementary Data 3 identified the immune effector complex, where the C1q complement machinery and fibrinogen tightly covary with immunoglobulins. This likely represents the protein corona^19^ or circulating immune complexes (CICs) that possess biophysical properties (size/density) overlapping with EVs. Similarly, Cluster 5, Supplementary Data 3 captured the hepatic secretome (CRP, SAA, Hemopexin), representing the soluble acute-phase response. Recognizing these modules is vital for biomarker discovery: a proteomic signature in a cancer patient may reflect the systemic hepatic response (Cluster 5) or paraneoplastic coagulation (Cluster 3/6, Supplementary Data 3) rather than tumor-derived vesicular cargo.

### Deconvoluting Cell Lysis from Active Secretion

Quality control in plasma proteomics has historically relied on single markers like hemoglobin. Our modular analysis provides a higher-resolution metric by distinguishing cytosolic lysis (Cluster 4, Supplementary Data 4: PKM, HBB, HBA1) from active secretion (Cluster 7, Supplementary Data 4: RAB27B, VASP).

The segregation of these clusters is biologically significant. It demonstrates that the release of cytosolic enzymes (lysis) and the shedding of membrane vesicles (secretion) are independent variance components in clinical samples. A sample can be high in platelet ectosomes (Cluster 5, Supplementary Data 4) without showing signs of lysis (Cluster 4, Supplementary Data 4), validating the physiological nature of the EV isolate. Conversely, the co-clustering of mitochondrial proteins *(ATP5F1B)* with the lysis module serves as a specific flag for cellular damage, as platelets and RBCs (the main EV sources) are either mitochondria-depleted or void.

In summary, we propose replacing binary EV vs. non-EV lists with a component-based reporting framework. Future plasma EV studies should quantify the relative abundance of at least these eight biological modules to establish the detailed characterization of the plasma EV samples. By deconvoluting the hepatic background and hematopoietic ectosomes from the endosomal signal, this atlas provides the necessary roadmap for discovering high-specificity biomarkers in liquid biopsy.

In summary, the analysis presented here challenges the prevailing view of the plasma EV proteome as a homogeneous entity. By deconvoluting the variance of 515 clinical samples, we demonstrate that the reproducible plasma EV isolate is likely a composite of at least eight distinct biological modules, only a subset of which are truly vesicular. Our data reveals that high-abundance proteins frequently cited as EV markers, including acute-phase reactants *(CRP, SAA)*, complement factors *(C1q, C3)*, and coagulation regulators, segregate into the Hepatic Secretome and Immune Effector clusters. These proteins represent a reproducible co-isolated plasma fraction driven by solubility and surface adsorption, distinct from vesicle biogenesis. Consequently, we argue that fluctuations in these markers should be interpreted as systemic liver responses or immune complex formation, rather than vesicular upregulation. Furthermore, we resolve the lineage attribution of canonical EV markers. The inextricable clustering of CD9 and ADAM10 with platelet and erythrocyte membranes (Cluster 7, Supplementary Data 3 and Cluster 5, Supplementary Data 4) suggests that these targets primarily monitor the total burden of blood cell ectosomes. In contrast, the distinct segregation of TSG101 and ALIX (Cluster 8) offers a specific metric for endosomal machinery that is independent of the bulk membrane background. Ultimately, this modular stratification transcends binary classifications of vesicular versus non-vesicular proteins. By mapping proteins to their specific biological source, whether the sub-membrane cortex, the lipid raft, or the soluble hepatic background, we provide a roadmap for high-resolution biomarker discovery that is robust to the inherent complexity of human plasma.

## Methods

### Systematic Search and Selection of Public Proteomics Datasets

#### Search Strategy

Keywords used to query the PRIDE repository: “plasma”, “extracellular vesicles”, “exosomes”, “serum” and “human”. Only data sets using label free quantitation and data dependent acquisition were maintained for the initial analysis. Data from the following accession were retrieved MSV000094307, PXD002668, PXD004420, PXD010404, PXD015283, PXD016293, PXD018301, PXD018337, PXD020629, PXD021718, PXD024072, PXD024216, PXD026483, PXD028515, PXD028885, PXD032767, PXD036735, PXD042379, PXD042422, and PXD0424222.

#### Inclusion

Datasets were retained if they contained >=3 raw files, human samples only, Label-free quantification compatible and using data dependent acquisition (DDA).

#### Exclusion criteria

Data set using data independent acquisition (DIA) and data sets with stable isotope based quantitation. DIA and stable isotope-labeled datasets are rare and technically challenging to compare directly with label-free DDA datasets.

This resulted in a data set of 515 LC-MSMS runs covering a broad range of plasma EV isolation methods. The methods cover differential centrifugation (DC), differential ultra-centrifugation (DUC), differential centrifugation followed by size exclusion chromatography (DC_SEC), Density gradient ultracentrifugation (DGUC), differential ultra-centrifugation followed by sucrose gradient (DUC_SUC), size exclusion chromatography (SEC), ExoRNeasy (Qiagen Italia, Milan, Italy), polymer-based precipitation followed by density gradient ultracentrifugation (PP_DGUC), polymer-based precipitation followed by size exclusion chromatography (PP_SEC), and PRotein Organic Solvent Precipitation (PROSPR).

### Unified Raw Data Processing (MaxQuant Workflow)

LC-MS data from previous studies based on human EV samples were downloaded from ProteomeXchange Consortium website^5,20-25^ (24th of March 2024). The data were analyzed by MaxQuant version (2.1) to ensure uniformity. The human UniProt FASTA database was used for database dependent search (3AUP000005640). Carbamidomethyl cysteine was specified as the fixed modification. Variable modifications included methionine oxidation, and N-terminal protein acetylation. Quantification was performed using label-free quantification (LFQ) (Fast LFQ enabled, and min. ratio count = 2) and intensity based quantification (iBAQ). The False Discovery Rate (FDR) threshold was set to 1% at peptide and protein levels. Reverse and contaminant matches were filtered out in R. To maintain the integrity of detection statistics, missing intensity values (NaN) were imputed with zero. This conservative approach was selected to strictly preserve the presence or absence information inherent in Data-Dependent Acquisition (DDA) mass spectrometry. Unlike probabilistic imputation methods (e.g., KNN or min-draw) that generate synthetic low-abundance values, zero-imputation ensures that the definition of the core plasma EV proteome remains restricted to proteins with genuine, reproducible spectral evidence. Consequently, proteins with sporadic detection profiles are penalized in mean-abundance calculations, ensuring that the downstream correlation analysis focuses exclusively on the robust, high-abundance fraction of the isolate. Finally, rank normalization was performed and the data filtered to maintain proteins detected in 50% of all LC-MS runs.

### Construction of High-Resolution Reference Proteomes for Non-Vesicular Components

To accurately deconvolute the plasma EV proteome, we established a compendium of high-depth reference datasets representing the primary non-vesicular sources of variance in plasma (e.g., platelets, erythrocytes, lipoproteins, and soluble immune complexes). This reference library was constructed through a multi-stage literature mining and manual curation pipeline designed to maximize specificity. First, we defined a panel of anchor proteins (Table 1) to serve as definitive, high-consensus markers for each non-vesicular biological compartment and EV markers. These anchors were selected based on their established utility in clinical diagnostics and cell biology as lineage-specific or compartment-specific indices (e.g., ITGA2B/CD41 for platelets, APOA1 for HDL, HBB for hemolysis). Using these anchor proteins as seeds, we mined high-resolution proteomic studies (studies referenced in Table 1) to compile a comprehensive associated proteome for each compartment which constitute proteins abundant in these compartments (Supplementary Data 1). This dataset encompasses all high-abundance proteins reported to functionally interact with or co-purify with the respective components. This list captures the broad interactome of plasma contaminants, including both integral component proteins and transiently associated binding partners. This list was then further processed. To ensure quantitative rigor in downstream analyses in results sections, we applied a strict manual curation filter to the associated proteome to generate a functional reference set (Supplementary Data 2). This step was critical to remove proteins that are merely associated (e.g., ubiquitous chaperones, non-specific matrix proteins) and retain only those with a clear functional annotation defining them as integral to the biological component. Criteria for inclusion in Supplementary Data 2 required obligate components of the macromolecular complex (e.g., *C1q* subunits for the complement complex). This curated Supplementary Data 2 served as the definitive gene set for the quantitative enrichment analyses presented in the results sections, ensuring that calculated abundance metrics reflected specific biological contamination burdens rather than generic protein noise.

### Statistical Analysis and Proteomic Variance Deconvolution

All statistical analyses and visualizations were performed within the R statistical programming environment (V4.2.2). To map the variance structure of the plasma EV proteome, rank-normalized iBAQ intensities were subjected to pairwise correlation analysis and unsupervised hierarchical clustering. Correlation and Clustering Methodology: To identify protein modules exhibiting consistent co-isolation patterns across the cohort, we calculated a pairwise Pearson correlation matrix for all proteins in the core and candidate datasets. Unsupervised agglomerative hierarchical clustering was performed on the correlation matrix using Euclidean distance as the dissimilarity metric and Ward’s minimum variance method (ward.D2) to minimize within-cluster variance. Heatmap Visualization and Cluster Definition: Clustered data were visualized using the pheatmap R package (V1.0.12). The optimal number of clusters *k* was determined by evaluating the dendrogram fusion heights (linkage distance). Inspection of the scree plot of linkage distances revealed a distinct plateau rather than a sharp elbow, indicating stable separation of the proteome into at least eight primary variance components. Consequently, the dendrogram was cut at *k*=8 using the cutree function (R stats package). Module Significance: Consistent with established benchmarks for plasma proteomics (Geyer *et al*., 2016^10^), correlation modules exhibiting a mean internal Pearson correlation coefficient of *r > 0*.*5* were defined as significant co-isolation units. This threshold distinguishes robust physical co-complexes and co-isolating particles from stochastic background noise.

## Acknowledgments

R.M. is supported by Fundaçâo para a Ciência e a Tecnologia (CEEC position, DOI: 10.54499/CEECIND/03906/2017/CP1421/CT0004). A.S.C. is supported by Fundaçâo para a Ciência e a Tecnologia (DOI 10.54499/DL57/2016/CP1457/CT0013). We receive funding from the European Union to advance EV research *(Horizon2020 GA n°* 101079264, EVCA *and Horizon Europe-SE2023/GA101183034)*. We acknowledge the COST Action CA20113 “PROTEOCURE” supported by COST (European Cooperation in Science and Technology). This work is funded by FEDER funds through the COMPETE 2020 Programme and National Funds through FCT—Portuguese Foundation for Science and Technology under the projects number PTDC/BTM-TEC/1746/2021. This work was supported by the Research Unit iNOVA4Health (UID/4462/2025) and by the Associated Laboratory LS4FUTURE (LA/P/0087/2020), all financially supported by Fundaçâo para a Ciência e Tecnologia / Ministério da Educaçâo, Ciência e Inovaçâo.

## Competing Interests/Declaration of Interest Statement

The authors declare no conflict of interest

## Author contribution

Conceptualization: RM; Resources RM; Data curation: ASC, JMF, HCB, RM; Software: RM; Formal analysis: ASC, RM; Validation: ASC, RM; Funding acquisition: ASC, RM; Visualization: RM; Writing - original draft: ASC, RM; Methodology: ASC, RM; Writing - review & editing: All authors.

---

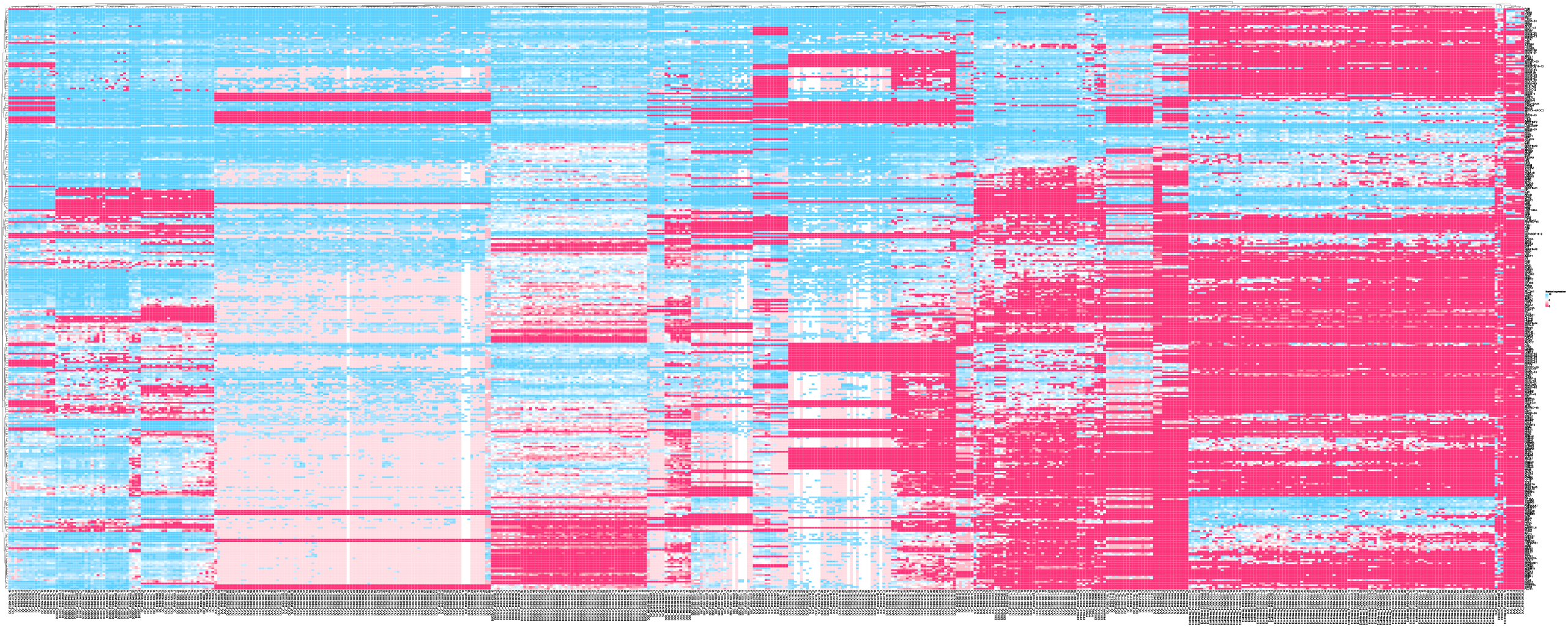

## Notes

### Competing Interest Statement

The authors have declared no competing interest.

