## Supplemental figures for "A Meta-Analysis of 515 Human Proteomes Defines the Core Composition and Contaminants of Plasma Extracellular Vesicles isolates"

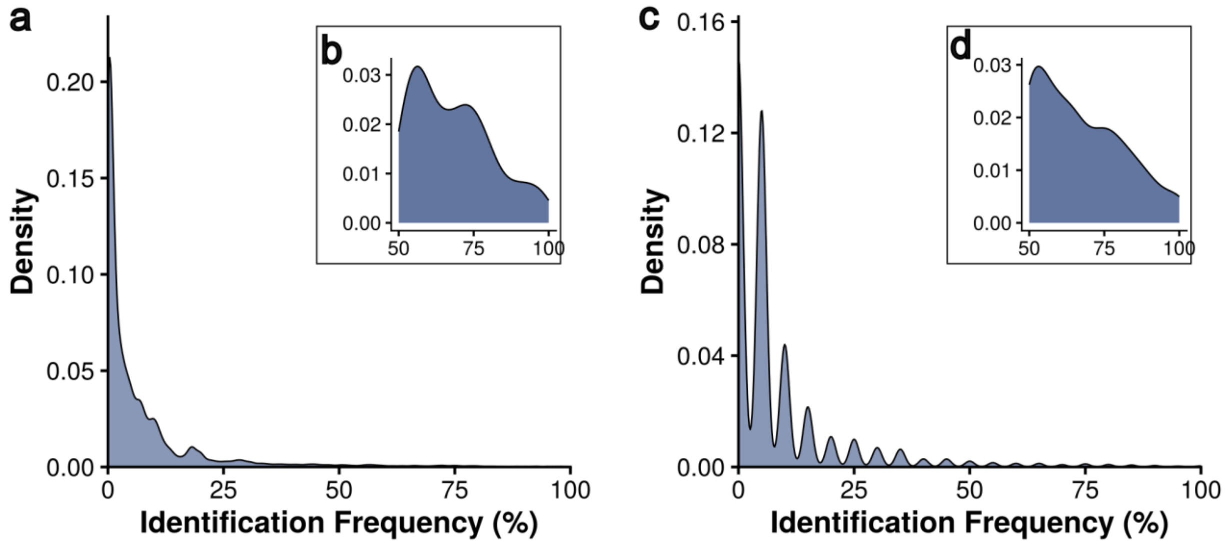


**Fig. S1 |** Identification frequency across **(a & b)** LC-MS runs and **(c & d)** studies.

**Fig. S2 | (Provide in separate file due to figure size). Heatmap with ranked protein expression values of proteins identified in more than 50% of all LC-MS runs.**

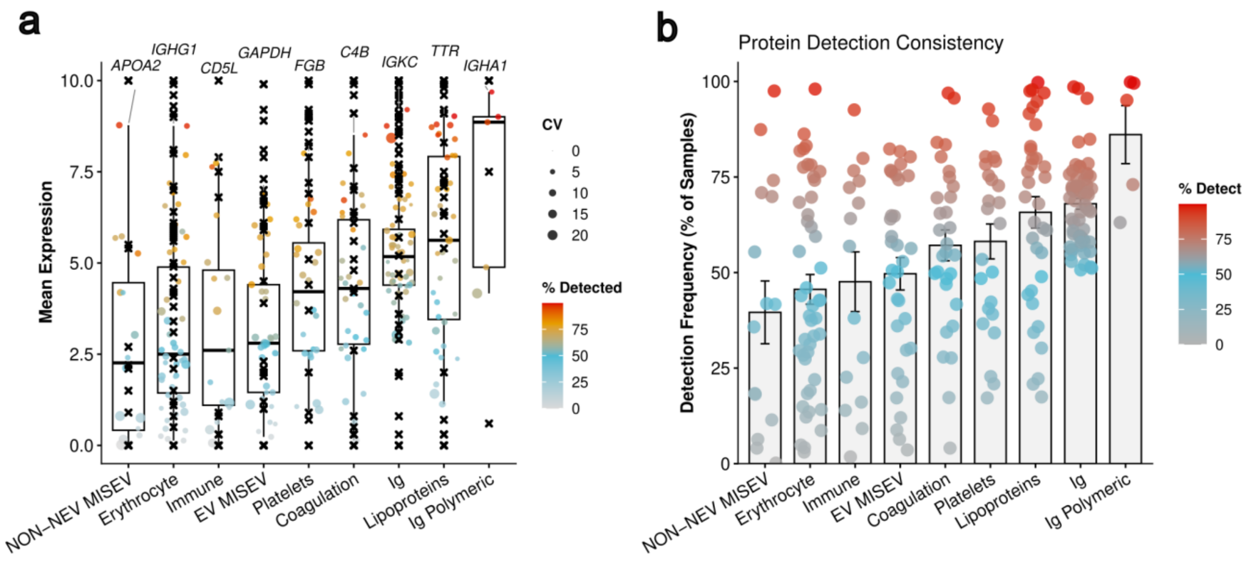


**Fig. S3 | Quantitative abundance and detection consistency of core EV markers in comparison with less strict non-EV markers signatures. a** Boxplots displaying the mean rank normalized iBAQ abundance of established EV markers (e.g., CD9, CD81) compared to common non-EV co-isolates (Lipoproteins, Erythrocyte proteins) across the entire dataset. The gene labels above each category is inidcating the most abundant protein in the core EV plasma proteome for the respective category. The central line represents the median, box limits indicate the 25th and 75th percentiles, and whiskers extend to 1.5× IQR. The crosses indicate abundance values from density gradient isolation of plasma EVs. **b** Detection frequency of each protein category, calculated as the percentage of the 515 LC-MS runs in which the marker was identified. Note that non-EV proteins are both abundant (**a**) and exhibit high detection consistency compared to core EV markers (**b**).
